# Effects of coil orientation on Motor Evoked Potentials from Orbicularis Oris and First Dorsal Interosseous

**DOI:** 10.1101/262261

**Authors:** Patti Adank, Dan Kennedy-Higgins, Gwijde Maegherman, Ricci Hannah, Helen Nuttall

**Affiliations:** Department of Speech, Hearing & Phonetic Sciences, University College London, UK; Sobell Department of Motor Neuroscience and Movement Disorders, Institute of Neurology, University College London, UK; Department of Psychology, Lancaster University, Lancaster, UK

**Keywords:** Transcranial Magnetic Stimulation, Motor Cortex, Facial muscle, Hand muscle, Motor Evoked Potentials, Coil orientation

## Abstract

**Objective:** This study aimed to characterise effects of coil orientation on the size of Motor Evoked Potentials (MEPs) from both sides of Orbicularis Oris (OO) and compare these effects with those reported for First Dorsal Interosseous (FDI), following stimulation to left lip and left hand Primary Motor Cortex.

**Methods:** Using a 70 mm figure-of-eight coil, we collected MEPs from eight different orientations while recording from contralateral and ipsilateral OO and FDI using a monophasic pulse.

**Results:** MEPs from OO were evoked consistently for six out of eight orientations for contralateral and ipsilateral sites. When latency and silent periods were taken into account, contralateral orientations 0°, 45°, 90°, and 315° were found to best elicit OO MEPs with a likely cortical origin. As expected, the largest FDI MEPs were recorded with an orientation of 45°, invoking a posterior-anterior (PA) current flow, from the contralateral location.

**Conclusion:** Orientations traditionally used for FDI were also found suitable for eliciting OO MEPs. Individuals vary more in their optimal coil orientation for eliciting MEPs from OO than for FDI. It is recommended that researchers iteratively probe several orientations when eliciting MEPs from OO. Care must be taken however because several orientations likely induced direct activation of facial muscles.

## 1. Introduction

The human motor system can be studied by recording Motor Evoked Potentials (MEPs) from skeletal muscles innervated by corticospinal or corticobulbar tracts (Cruccu et al., 1990, Rothwell et al., 1999, Rossini et al., 2015). MEPs are crucial in characterising motor system function in a variety of tasks (Pascual-Leone et al., 2000), including its interaction with a variety of drugs in healthy subjects and patients with damage to the central nervous system (Rossini et al., 2015). Several consensus papers prescribe procedures for standardized MEP collection, specific to corticospinal (Rossini et al., 2015) and corticobulbar (Groppa et al., 2012) systems. One well-known feature of hand muscle MEPs, innervated by the corticospinal tract, is their sensitivity to coil orientation and thus orientation of the induced current in the brain (Brasil-Neto et al., 1992, Mills et al., 1992, Werhahn et al., 1994). The recommended coil orientation for collecting MEPs from the hand is at an angle of 45° (cf. Figure 1) with respect to the sagittal plane, which induces a posterior-anterior (PA) current flow approximately perpendicular to the anterior wall of the central sulcus, as this evokes MEPs with the lowest stimulus intensities (Rossini et al., 2015).

**Figure 1.**
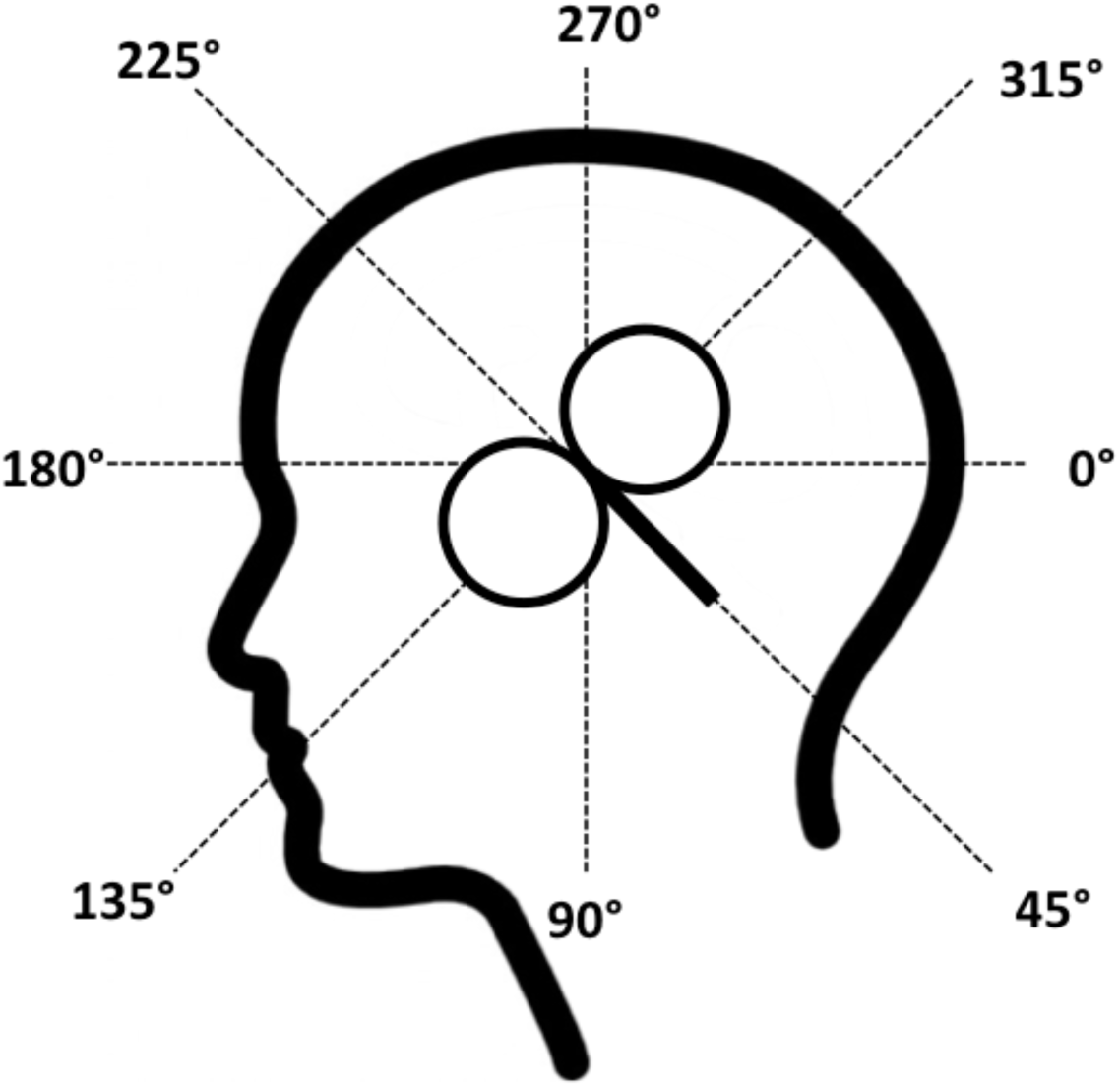
Eight coil orientations used in the lip and hand conditions. The intersection of the lines was placed on the subject’s hot spot for lip or hand M1 and the coil handle was aimed towards the angle tested (here: 45°).

**Figure 2.**
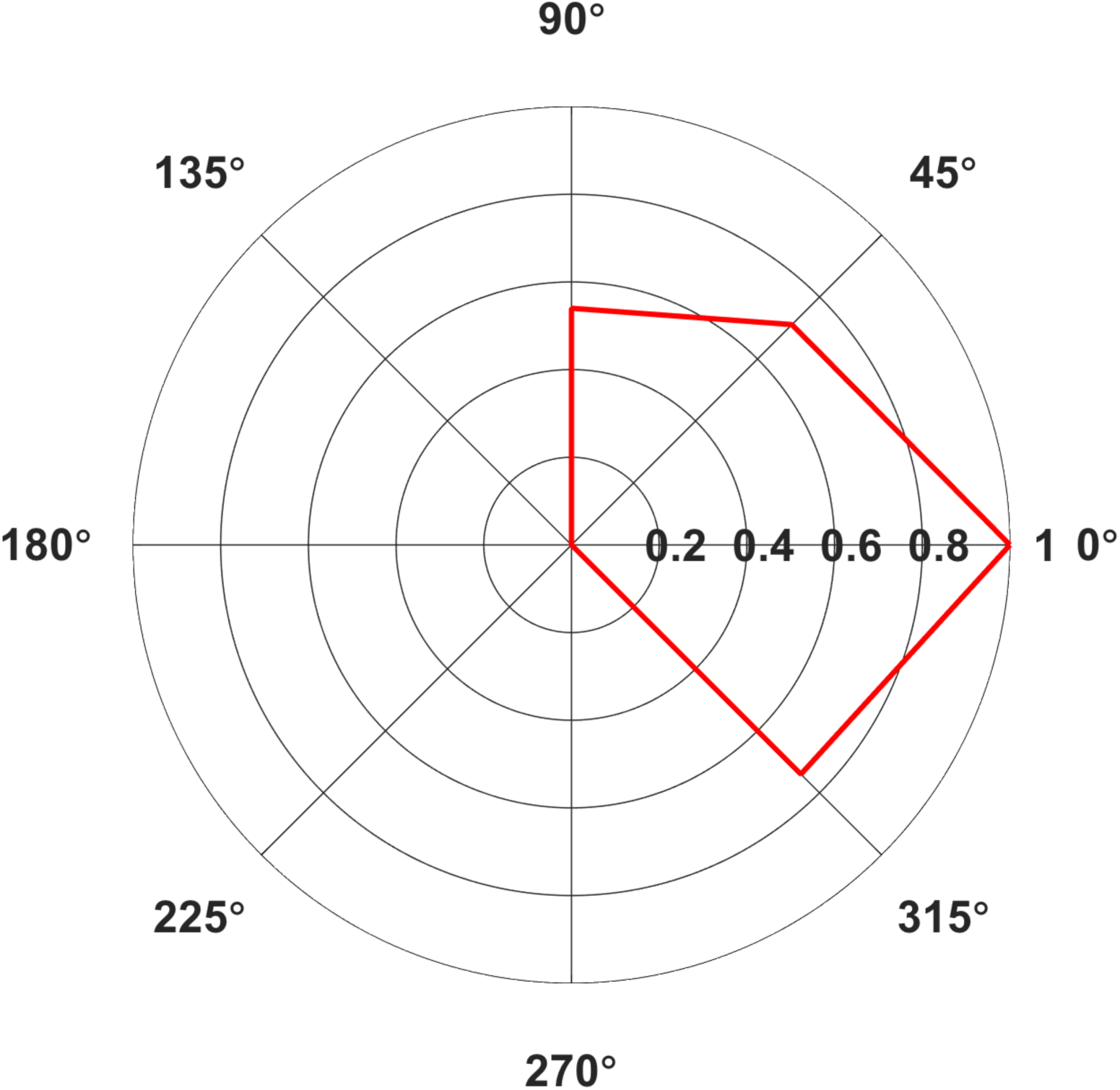
Polar plot of average Area Under the Curve (AUC) elicited from contralateral (A) Orbicularis Oris (OO) in mV·ms for Middle Latency MEPs only. Only average values with >5 contributing subjects are included. Values normalised relative to the largest value, set to 1.

When stimulating the corticospinal tract, two main patterns can be observed. First, low intensity PA currents preferentially evoke short-latency responses with the lowest thresholds, whilst anterior-posterior (AP) directed currents have a higher thresholds and preferentially evoke longer latency responses (Day et al., 1989, Sakai et al., 1997, Hannah et al., 2017b). These differences are thought to reflect the recruitment of different sets of excitatory synaptic inputs to corticospinal axons, with different physiological (Hanajima et al., 1998, Hannah et al., 2017b) and behavioural properties (Hamada et al., 2014, Hannah et al., 2017a). Second, latero-medial (LM) oriented currents preferentially evoke the shortest latency responses likely resulting from direct activation of corticospinal axons, i.e. non-synaptic activation (D-waves), and thus are sub-optimal for evaluating changes in cortical excitability since they effectively bypass cortical synapses. However, while the optimal coil orientation – and resulting induced current direction – for eliciting MEPs from muscles innervated by the corticospinal tract is fairly well-established, recommendations are less clear for facial muscles innervated through the corticobulbar tract. A handful of studies report on the effect of coil orientation on MEPs elicited for various facial muscles, including Nasalis (Dubach et al., 2004), Masseter (Guggisberg et al., 2001), Depressor Anguli Oris and Depressor Labii Inferioris (Rodel et al., 1999), and muscles in the tongue (Murdoch et al., 2013). For instance, Guggisberg et al. reported largest MEP peak-to-peak (p-p) amplitudes at 120° and 300°For Masseter, orientations that are approximately perpendicular to the 45° orientation found to be optimal for hand muscles. Murdoch et al. found that optimal orientations extended between 135° and 225° across subjects for tongue muscles. Thus, it seems clear that the optimal angle for the hand area does not apply to the facial muscles. Furthermore, given the uncertainty concerning the optimal orientation across facial muscles, a detailed exploration of the optimal angle for evoking MEPs seems necessary prior to investigating facial muscles such as the Orbicularis Oris (OO) for which there are no current guidelines.

The OO muscle is relevant to both clinical research, e.g., in Bell’s Palsy (Schriefer et al., 1988, Meyer et al., 1994), and cognitive neuroscience studies where OO MEPs have been recorded to investigate changes in motor cortex activity during speech perception (Fadiga et al., 2002, Roy et al., 2008, Murakami et al., 2011, Swaminathan et al., 2013, Nuttall et al., 2016, Nuttall et al., 2017). These studies induced a PA current flow by adopting the standard 45o downward pointing orientation recommended for hand MEP acquisition, despite the fact it has not been explicitly verified that the 45o orientation used for hand MEPs is also suitable for OO. From a practical point of view, using an optimised coil orientation for OO will allow clinical and cognitive neuroscience researchers to use lower stimulus intensities, thus potentially reducing subject dropout due to discomfort. Importantly, it may also help avoid the potential for direct activation of the corticobulbar axons (D-waves), which might otherwise confound the proper interrogation of the corticobulbar pathway.

Given the similar organisation of the lip and hand motor cortex, a similar anisotropy of lip motor cortex responses to TMS might be assumed. Furthermore, it seems plausible that the optimal coil orientations for lip and hand are similar given their similar location in the lip of pre-central gyrus / anterior bank of the central sulcus (Penfield et al., 1937) and the broadly similar orientation of the sulcus at each point. However, unlike in hand muscles, the location of the coil close to cranial nerves when eliciting facial MEPs lends to the possibility of directly activating motor nerves supplying facial muscles, to evoke M-waves, which could contaminate measurement of MEP amplitudes and latencies. The likelihood of directly activating facial muscles could depend on proximity of the coil to the nerve, and therefore on coil orientation. Dubach et al. (2004) reported direct innervation of ipsilateral and contralateral Nasalis muscle using surface electrodes; they report short latency responses (<7.5 ms) with a likely peripheral origin. These short-latency-responses were most commonly elicited for coil orientations 120° and 165°, i.e., orientations with the coil handle facing in an antero-ventral direction (i.e., directly over the subject’s temple), inducing approximately LM or AP currents. Given the proximity of the different facial regions in the cortex, we expected that responses evoked by TMS in OO might also consist of both peripherally- (M-waves) and cortically-evoked (MEPs) responses, and that the presence of M-waves would vary according to coil angle.

This study aimed to determine the effect of the direction of the induced current on the size and morphology of contra- and ipsilateral MEPs of the OO muscle by systematic manipulation of the coil orientation used to evoke MEPs. We measured MEPs evoked from OO and First Dorsal Interosseous (FDI), to enable comparison of OO results with the well-documented effect of current direction and coil orientation on FDI (Werhahn et al., 1994, Balslev et al., 2007), following stimulation of left lip and hand M1, respectively. We measured MEPs for eight coil orientations following stimulation to the left hemisphere only. We also examined the possibility that direct nerve innervation for lip muscles: (i) by measuring OO MEPs from contra- and ipsilateral sites of OO, since M-waves would only be expected to be present on the side ipsilateral to stimulation; and (ii) by measuring the onset latencies of responses as well as the presence of a cortical silent period to determine their likely origin (i.e. cortical and synaptic versus peripheral and non-synaptic).

## 2. Method

### 2.1 Subjects

We tested sixteen subjects (seven males; average age: 30 years 2 months ± SD 6 years 11 months; age range: 24–47 years, 10RH, 6LH). Handedness was established via the Edinburgh handedness inventory (Oldfield, 1971). Data from one right-handed male subject were discarded due to a technical error during data collection. Subjects presented no TMS contraindications, and did not report any (history of) neurologic/psychiatric disease, or that they were under the effect of neuroactive drugs. All subjects had a minimum high school-level education, with the majority currently studying at University level. They were asked to not consume caffeine-containing drinks before the start of the experiment. All subjects were tested in the morning. Experiments were undertaken with the understanding and written consent of each subjects, according to University College London Research Ethics Committee (UREC).

### 2.2 Transcranial magnetic stimulation

Monophasic single TMS pulses were generated by a Magstim 200^2^ unit and delivered by a 70 mm diameter figure-of-eight coil, connected through a BiStim2 module (Magstim, Dyfed, UK) set to simultaneous discharge mode (inter-pulse spacing of 0 ms). The coil was placed tangential to the skull such that the induced current flowed from posterior to anterior under the junction of the two wings of the figure-of-eight coil. The lip and hand areas of M1 was found using the functional ‘hot spot’ localization method, whereby application of TMS elicits an MEP from the contralateral muscle. We located the hand area by asking subjects to press their index finger upwards against the desk surface to activate FDI. The lip area ‘hot spot’ was identified by moving the coil ventrally and slightly anterior until consistent MEPs were observed in the contralateral lip muscle, and the aMT identified (Möttönen et al., 2014). The coil position and orientation was adjusted in small movements to ascertain the location on the scalp at which the largest and most consistent MEPs are elicited. This location was then marked on a cap and active motor threshold (aMT) determined, which constitutes the intensity at which TMS pulses elicited five out of 10 MEPs with an amplitude of at least 200*µ*V for OO, and at least 500*µ*V for FDI at 20% of maximal contraction. The intensity of the stimulator was then set to 120% of aMT for the stimulations applied during the experiment. The mean aMT used to elicit OO MEPs was 48.0% (± 5.6%), and 39.4% (± 6.4%) of the maximum possible intensity for FDI MEPs. Testing occurred at 120% of aMT; 57.6% (± 6.8%) for OO and at 47.4% (± 7.6%) for FDI.

### 2.3 Electromyography

Electromyographic (EMG) activity was recorded from lip and hand areas using surface electrodes (Ag/AgCl; 10-mm diameter) in a non-Faraday caged, double-walled sound-attenuating booth. For the lips, electrodes were attached to OO on the both sides of the mouth, on the upper lip, approximately 5 mm from the vermillion border, orientated horizontally, in a bipolar, belly-belly montage, with electrodes placed at the left and right temples serving as a common ground. To stabilize background EMG activity, subjects were trained for approximately five minutes to produce a constant level of contraction (approximately 20% of maximum voluntary contraction) of the lip muscles by pursing, which was verified via visual feedback of the on-going EMG signal. For the recording of hand EMG, electrodes were attached in a tendon-belly montage with the active electrode placed on both FDI muscles, the reference electrode on the tendon of the same muscle, and a ground electrode on each wrist. Subjects were also trained to maintain a constant level of contraction of this muscle during the experimental recordings. Contraction of the lip and hand muscles also facilitates a lower motor threshold relative to when the muscle is at rest, enabling the use of lower levels of stimulation during the experiment. It is not straightforward to elicit MEPs from OO in the relaxed muscle because of the relatively high threshold. The raw EMG signal was amplified by a factor of 1000, band-pass filtered between 100-2000 Hz, and sampled at 5000 Hz online using a 1902 amplifier (Cambridge Electronic Design, Cambridge), and analog-to-digital converted using a Micro1401-3 unit (Cambridge Electronic Design, Cambridge). Continuous data were acquired and recorded using Spike2 software (version 8, Cambridge Electronic Design, Cambridge).

### 2.4 Procedure

Following recommendation from Groppa et al. (2012) for exploratory non-clinical studies, we delivered TMS pulses to eight orientations: 0°, 45°, 90°, 135°, 180°, 225°, 270°, 315° (Figure 1) in each subject for both sides of OO after identification of the hotspot and motor threshold for left lip M1. In a separate block, we delivered TMS pulses from the same orientations after stimulation of the hot spot for left hand M1 from left and right FDI. We therefore aimed to collect 480 MEPs over two channels per subject per muscle (960 in total). Subjects maintained 20% of maximal voluntary contraction in both hands, to make the results of the ipsilateral recordings comparable with both lip channels (it is not straightforward to relax only one side of OO). We counterbalanced the order in which hand and lip MEPs were collected across subjects. Within a lip or hand block, the coil rotation procedure was as follows. The subject wore an unmarked EEG cap. After localising the hot spots for lip and hand M1, we attached a pre-constructed lattice made of adhesive tape aligned in a starburst pattern made of tape to the EEG cap. This lattice contained four intersecting guiding lines as in Figure 1. The intersection of the lines was placed on the hot spot for lip or hand M1, ensuring that the line between 180° and 0° in Figure 1 was approximately level with the parasagittal plane. The same lattice was used for all subjects. For each orientation, the handle of the coil was parallel with the guiding line for the target angle and the intersection between the two wings of the coil was on the intersection and on the hot spot. Figure 1 shows the position of the coil’s wings and handle for obtaining MEPs for the 45° angle with a PA current direction. The coil aligned on the subject’s head using the tape guide, and 30 MEPs per orientation were collected. We used a relatively high number of MEPs compared to previous studies (Guggisberg et al., 2001, Dubach et al., 2004). A low number of trials per subject is associated with decreased statistical power and with inflated variability and noise. It has therefore been suggested to record at least 20–30 MEPs per condition in basic and clinical settings (Schmidt et al., 2009, Cuypers et al., 2014, Goldsworthy et al., 2016). After completing testing for a single orientation, subjects were asked to report muscle twitches and their level of comfort (1–7, 7 high comfort). The order for all eight angles was randomised as across subjects and the same order was used across lip and hand blocks. The duration of the session was between 2.5 and 3 hours.

### 2.5 Data analysis

Individual EMG sweeps starting 40 ms before the TMS pulse and ending 1000 ms post-stimulation were exported offline from the recording software into Matlab, where mean MEPs were calculated for the two channels per TMS target, orientation, and subject. Individual averages were rectified and the integrated area under the curve (AUC) of the rectified EMG signal of each individual mean MEP was calculated as millivolts over millisecond (mV·ms). We chose to measure AUC instead of p-p – a measure commonly used for hand MEPs – as OO MEPs tend to consist of multiple peaks, in contrast with hand MEPs, which tend to consist of two successive midline deflections (peak and trough) (Adank et al., 2016). It is not straightforward to measure p-p amplitudes when successive peaks are present, and the amplitude of lip MEPs can be underestimated if p-p is used as successive peaks in an OO MEP complex would be excluded from the final amplitude measurement. For lip MEPs, AUC was automatically computed from 8 to 35 ms post-TMS, and for hand AUC was computed from 13 to 40 ms post-TMS.

To confirm that the AUCs reflected cortically generated MEPs, and not the activity resulting from direct innervations of facial nerves, we analysed the data taking into consideration latency, CSP, and reported muscles twitches. The analysis of latencies and CSPs was conducted by hand, in a separate process from the AUC analysis (which was conducted using Matlab scripts). Short latencies suggest that the MEPs originated via direct innervation of facial muscles (cf. Dubach et al. 2004). We split the OO data into short latency (<7.5 ms) and middle latency (≥7.5 ms) responses, following Dubach et al. MEPS evoked during contraction are generally followed by a period of silence, i.e. interruption of volitional EMG, before normal voluntary activity resumes. Therefore the presence of a silent period might be useful in determining the nature of the response. The TMS-induced CSP is thought to have a cortical origin and result from intracortical inhibition, while silent periods following direct electrical stimulation of peripheral muscles innervated by the corticospinal tract are linked to peripheral inhibition of spinal motoneurons (Wilson et al., 1993). However, electrical stimulation of muscles in the corticospinal tract may not to result in silent periods (Cruccu et al., 1997, Katayama et al., 2001). We measured silent periods to give an indication about the likely origin of the evoked MEPs. However, it should be noted that not all cortically evoked MEPs will necessarily evoke a clear CSP, and that the lack of a CSP can therefore not be used as measure for excluding a likely cortical origin. Finally, we asked subjects to report whether they experienced muscle twitches for each orientation to attempt to provide further information about the potential origin of MEPs. If eliciting MEPs for a specific orientation results in MEPs with a short latency, no CSP, and subject report a high proportion of muscle twitches in face or jaw, then the MEP is likely to be the result of direct innervation of the facial nerves. Conversely, it is more plausible that the MEP had a cortical origin if it was middle latency, showed a clear CSP, and subjects did not report a high proportion of facial twitches for the associated orientation. The average MEPs collected from ipsi- or contralateral muscles and for the eight orientation were compared using a non-parametric ANOVA that allowed for missing values, the Skillings-Mack test (Chatfield et al., 2009), for both effectors (hand or lip) separately.

## 3. Results

### 3.1 Lip

Mean AUCs for OO are reported in Table I and latencies and CSPs are listed in Tables II and III, respectively. MEPs from OO for 135° were not recorded for a male left-handed subject due to a technical error. Seventeen of 240 average MEPs were classified as outliers (>1.5 the interquartile range) and excluded from further analysis.

**Table I:** Average Area Under the Curve (AUC) in mV·ms, plus standard deviations (SD), and number of subjects (N) contributing to the average, for orbicularis Oris (OO) and First Dorsal Interosseous (FDI) muscles per coil orientation.

|  | Contralateral |  |  | Ipsilateral |  |  |
| --- | --- | --- | --- | --- | --- | --- |
| OO | AUC | SD | N | AUC | SD | N |
| 0° | 8.6 | 7.0 | 15 | 3.3 | 1.1 | 12 |
| 45° | 8.2 | 5.4 | 14 | 3.5 | 1.2 | 12 |
| 90° | 6.3 | 3.5 | 15 | 3.4 | 2.0 | 14 |
| 135° | 6.8 | 3.2 | 15 | 6.2 | 5.7 | 13 |
| 180° | 6.3 | 3.6 | 15 | 9.5 | 8.7 | 14 |
| 225° | 2.7 | 1.4 | 15 | 2.2 | 0.7 | 15 |
| 270° | 2.9 | 1.5 | 14 | 2.0 | 0.9 | 13 |
| 315° | 7.2 | 5.7 | 15 | 3.3 | 1.7 | 11 |
| FDI | AUC | SD | N | AUC | SD | N |
| 0° | 21.7 | 15.4 | 15 | 2.5 | 1.3 | 14 |
| 45° | 36.3 | 17.2 | 15 | 2.8 | 1.8 | 15 |
| 90° | 10.9 | 9.2 | 13 | 2.0 | 1.3 | 14 |
| 135° | 5.9 | 6.1 | 15 | 2.9 | 1.7 | 15 |
| 180° | 5.9 | 7.7 | 15 | 2.7 | 1.3 | 15 |
| 225° | 4.2 | 1.5 | 12 | 2.3 | 1.2 | 13 |
| 270° | 4.8 | 3.5 | 13 | 2.8 | 1.3 | 14 |
| 315° | 5.7 | 5.0 | 14 | 2.7 | 1.1 | 15 |

**Table II:** Average response latency duration in milliseconds, plus standard deviations (SD), and number of subjects (N) contributing to the average, for orbicularis Oris (OO) and First Dorsal Interosseous (FDI) muscles per coil orientation.

|  | Contralateral |  |  | Ipsilateral |  |  |
| --- | --- | --- | --- | --- | --- | --- |
| OO | Latency | SD | N | Latency | SD | N |
| 0° | 9.6 | 1.1 | 6 | - | - | - |
| 45° | 9.7 | 1.2 | 7 | 6.3 | - | 1 |
| 90° | 8.9 | 0.7 | 6 | 8.3 | 2.8 | 2 |
| 135° | 7.4 | 1.4 | 9 | 6.9 | 1.7 | 4 |
| 180° | 6.7 | 0.8 | 8 | 6.6 | 0.7 | 7 |
| 225° | - | - | - | - | - | - |
| 270° | - | - | - | - | - | - |
| 315° | 9.1 | 2.5 | 6.0 | - | - | - |
| FDI | Latency | SD | N | Latency | SD | N |
| 0° | 20.8 | 2.1 | 10 | - | - | - |
| 45° | 20.8 | 1.9 | 14 | 6.3 | - | 1 |
| 90° | 20.8 | 1.5 | 3 | 8.3 | 2.8 | 2 |
| 135° | 21.5 | - | 1 | 6.9 | 1.7 | 4 |
| 180° | 27.0 | - | 1 | 6.6 | 0.7 | 7 |
| 225° | - | - | - | - | - | - |
| 270° | - | - | - | - | - | - |
| 315° | 22.0 | 3.5 | 2 | - | - | - |

**Table III:** Average Cortical Silent Period (CSP) duration in milliseconds, plus standard deviations (SD), and number of subjects (N) contributing to the average, for orbicularis Oris (OO) and First Dorsal Interosseous (FDI) muscles per coil orientation. No CSPs were measured for any of the right hemisphere orientations for either muscle.

| Orientation | OO Contralateral |  |  | FDI Contralateral |  |  |
| --- | --- | --- | --- | --- | --- | --- |
|  | CSP | SD | N | CSP | SD | N |
| 0° | 20.0 | 3.7 | 4 | 32.0 | 18.1 | 6 |
| 45° | 16.2 | 2.9 | 5 | 43.9 | 28.3 | 8 |
| 90° | 15.0 | 1.4 | 2 | - | - | - |
| 135° | - | - | - | - | - | - |
| 180° | - | - | - | 37.0 | - | 1 |
| 225° | - | - | - | - | - | - |
| 270° | - | - | - | - | - | - |
| 315° | 18.5 | 11.6 | 4 | 14.5 | 2.1 | 2 |

Latencies (Table II) were measured for at least 5 out of 15 subjects for contralateral OO in 0°, 45°, 90°, 180°, and 315°, in ipsilateral OO for 180°. Middle latency responses were recorded for contralateral 0°, 45°, 90°, and 315° only. CSPs (Table III) were measured in contralateral only for 5 out of 15 subjects for 45°. Short latencies were found for 135° and 180°, no CSPs could be measured for either of these orientations in contra nor ipsilateral OO. Figure 3 shows raw MEPs for one subject collected from contra- and ipsilateral orientations to illustrate effects of coil orientation on latency and CSP. Note the short latency for response collected from 180°Contralateral and 135° ipsilateral orientations compared to the latency for 45°Contralateral. Moreover, for 45°Contralateral, a clear CSP was detected, while there was no clear interruption of voluntary EMG following the evoked response for the 135°Contralateral and 0° and 180° ipsilateral orientations. Moreover, subjects reported muscles twitches on 50 occasions: in jaw (30), eye (11), forehead (2), face (2), neck (3), lip (1), or nose (1). Note that the presence of a muscle twitch does not automatically imply direct nerve activation, but that muscle twitches could also result from a twitch in the target muscle. The majority of facial twitches were reported for 180° (11), followed by 0° (9), 270° (7), and 45° (6). Note that TMS to lip M1 generally does not result in noticeable twitches in OO, and it was reported only once throughout the entire experiment. The number of reported twitches for lip was higher than for the hand condition (15), presumably as on average higher stimulator output was used in the lip condition (48% versus 38% for the hand condition). Also, a more ventral, and for most subjects also more anterior, placement of the coil was used to target lip M1, and the coil was therefore closer to superior branches of the facial nerve (i.e., the temporal and zygomatic branches).

**Figure 3.**
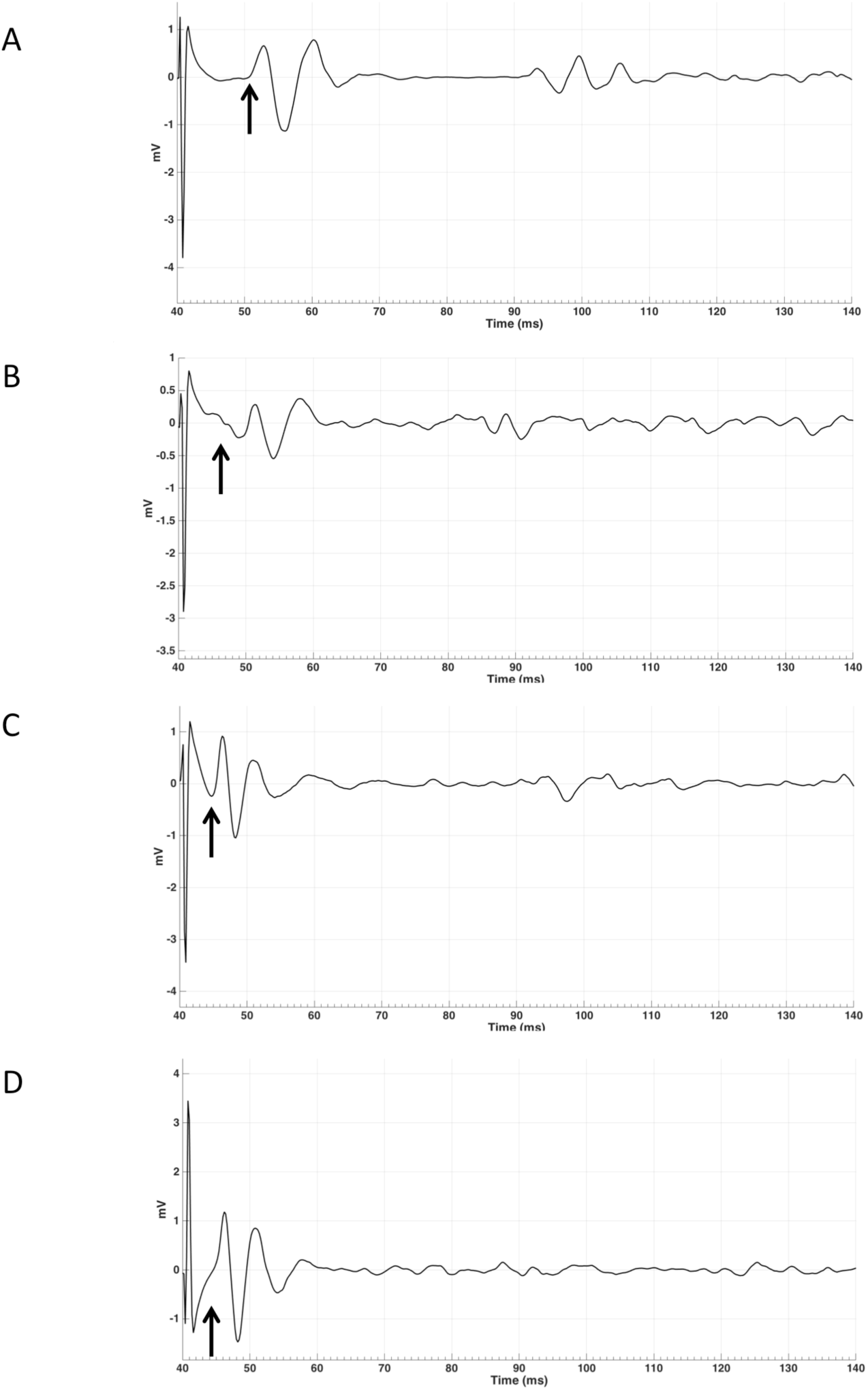
MEP EMG traces for a single MEP for subject 9 for four orientations: contralateral 45 (A), contralateral 135 (B), ipsilateral 0 (C), and 180 (D) in mV. Arrows indicates approximate start of MEP, measured from TMS pulse at 40 ms into the trial.

Based on the measurements of latencies and subject reports of facial twitches, it appears that MEPs with a cortical (synaptic) origin were plausibly invoked predominantly in contralateral sites with the coil handle positioned at 315°, 0°, 45° and 90° (Figure 1), and in two subjects in 90° ipsilaterally. MEPs generated in all ipsilateral orientations, potentially with the exception of 90°, and contralateral orientations 135°, 180°, 225°, and 270° likely had a peripheral origin (but we cannot exclude the possibility that these responses were the result of direct activations of contralateral corticobulbar) and may have resulted from direct stimulation of facial motor nerves. We included the averages for contralateral orientations 0°, 45°, 90°, and 315° in the Skillings-Mack test (contralateral 135° was not included as the test does not allow levels with fewer than three observations) and found no difference in AUC (t(3)=5.0589, p=0.1675). This indicated that these four orientations resulted in comparable AUCs. However, it seems like contralateral 0° and 45° are optimal for evoking OO MEPs, as this orientation resulted in the highest proportion of middle latency MEPs that also showed CSPs (Tables II and III).

### 3.2 Hand

Mean AUC results for FDI are reported in Table I, and latencies and CSPs are listed in Tables II and III, respectively. One subject erroneously did not contract her right hand during collection of bilateral FDI MEPs. Fifteen out of 240 average MEPs were classified as outliers and were excluded from further analysis. Subjects reported muscles twitches in their hand (7), jaw (2), eye (2), face (2), nose (1), or neck (1) on 15 occasions. Facial twitches were reported most often for 135° (4), followed by 180° (3). We compared the average AUC values between contralateral orientations 0° and 45° in the Skillings-Mack test and found a significant difference in AUC (t(1)=4, p=0.0455), with larger AUCs for 45° than for 0° (Figure 4).

**Figure 4.**
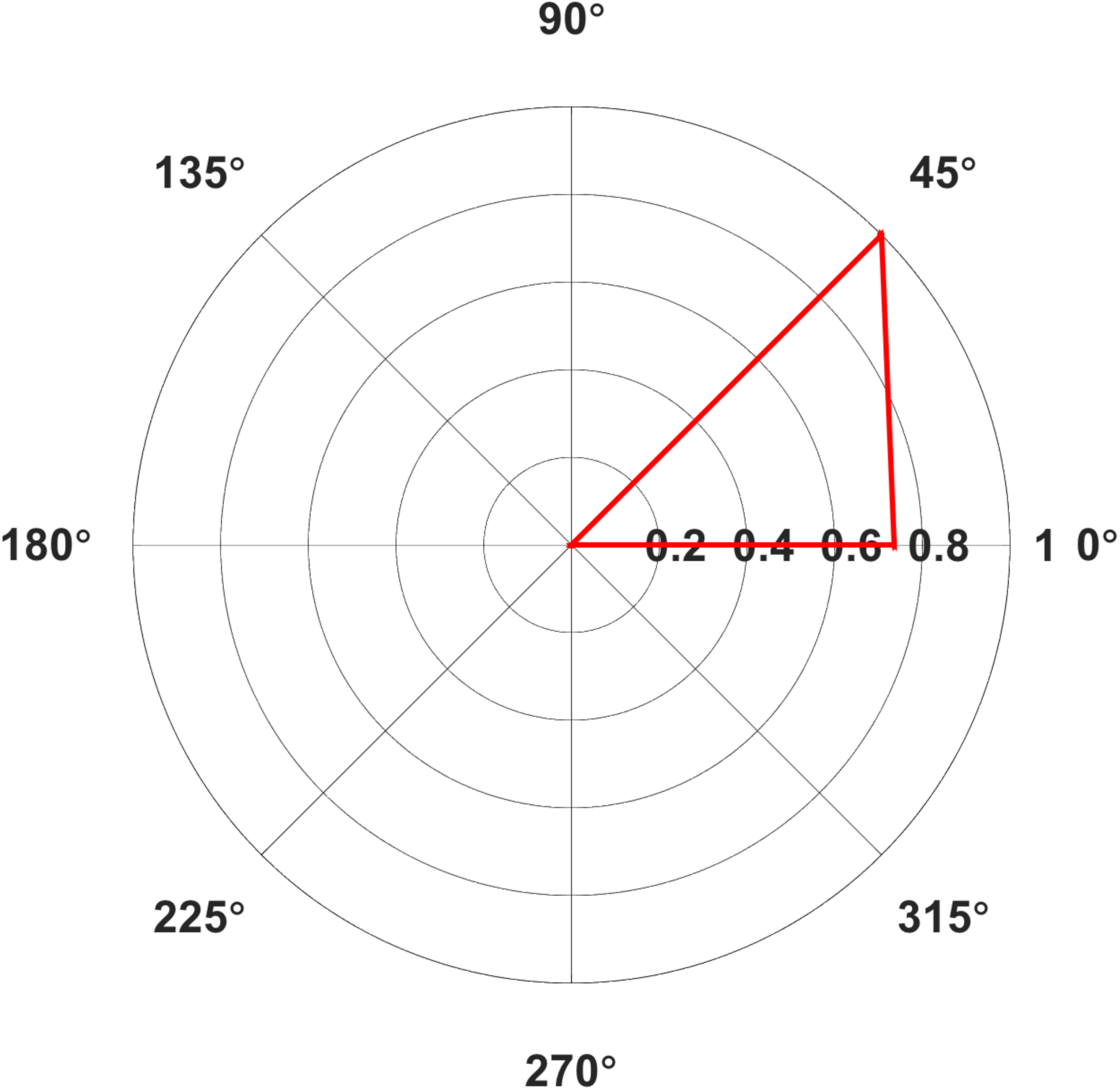
Polar plot of average Area Under the Curve (AUC) elicited from contralateral (A) First Dorsal Interosseous (FDI) in mV·ms for Middle Latency MEPs only. Only average values with >5 contributing subjects are included. Values normalised relative to the largest value, set to 1.

## 4. Discussion

Our results replicated previous findings by showing that 45° was the optimal orientation for FDI MEPs. Middle latency (~20 ms) responses associated with a clear CSP measurement were found only for contralateral 45° and 0°. MEPs recorded from ipsilateral FDI had short latencies (<10 ms) and no CSP was measured for any of the recording from ipsilateral FDI. The results for FDI replicate findings previously reported for studies investigating the effect of coil orientation (Pascual-Leone et al., 1994, Werhahn et al., 1994, Balslev et al., 2007).

For OO, after controlling for the presence of potential non-cortical responses, it seemed that MEPs could be evoked across a broader range of contralateral angles, 0°, 45°, 90°, and 315°, and these were also the only orientations that generated middle latency MEPs. Ipsilateral responses were rare and tended to include short-latency responses that are most likely due to direct activation of facial nerves. Specifically, the OO results showed largest MEP amplitudes for contralateral. It is not clear why the range of orientations (and associated currents) is wider for OO than for FDI. Speculatively, this difference between OO and FDI may be due to differences in the specific anatomy of the cortex, leading to a less homogenous orientation of neurons in the lip area compared to the hand area. Moreover, the optimal orientation for OO varies across individuals and that optimal MEPs for an individual might be collected at 0°, 45°, 90°, or 315°. As was the case for previous results on the effect of coil orientation on elicitation of MEPs in other facial muscles (Rodel et al., 1999, Pilurzi et al., 2013), we measured ipsilateral OO MEPS in several subjects, although the low amplitudes and short latencies found for the majority of subjects indicate that most were likely M-waves. Rödel et al. report ipsilateral MEPs in OO for all their subjects and also report slightly larger MEP amplitudes for ipsilateral MEPs for a subset of their tested coil positions relative to vertex, while Pilurzi et al. report ipsilateral MEPs for 14 out of 18 subjects for the Depressor Anguli Oris muscle. Moreover, (Triggs et al., 2005) collected contra and ipsilateral MEPs from OO after stimulating left and right M1. Ipsilateral MEPs were larger than contralateral MEPs after TMS to left and right lip M1 in a subset of Triggs et al.’s subjects (14 of 42), in line with our results for OO from several ipsilateral orientations (Table I). Ipsilateral MEPs in OO may originate from cortical sources from I-waves generated by pyramidal neurons in M1, or through direct innervation of the facial nerves, as discussed earlier. Even though OO is the only median facial muscle, it does not seem likely that the recorded MEPs were due to action potentials travelling across muscle fibers between both sides of the lips. When muscular fibres in OO were directly stimulated innervated via the facial nerve ipsilaterally have been reported to cross the midline for only a few millimetres (Trojaborg, 1977). Midline crossing of motor axons in facial nerves can occur in pathological conditions such as complete unilateral facial palsy (Gilhuis et al., 2001). The subjects reported the highest proportion of facial (jaw) twitches for 180°, an orientation in which the coil handle is positioned (Figure 1) so that it seems feasible to directly stimulate upper branches of the facial nerve. However, lower face twitches were also reported for other orientations (0°, 270°, 45°), so it seems implausible that orientation can be directly linked to facial nerve innervation. Our data does not allow for conclusive elimination of the possibility that ipsilateral MEPs were evoked by direct stimulation of the ipsilateral facial nerve. Therefore, to clarify the origin of ipsilateral MEPs, and of MEPs evoked from contralateral orientations 135°, 180°, and 270° it would be useful, for example, to employ a paired-pulse protocol such as SICI (Short-Interval Intra-cortical Inhibition) (Kujirai et al., 1993), to clarify the neural origin of MEPs for different current directions. SICI is present in lower facial muscles, as demonstrated by (Pilurzi et al., 2013).

We suggest taking an individualised approach to determining the optimal rotation for lip muscles. A comparable approach was suggested by Murdoch et al. for tongue muscles. They proposed to probe several coil orientations until one is found that consistently produces the largest MEPs. They suggest to initially examine “a range of directions extending from 315° to 135° or rather dorsal-caudal through to ventral-rostral”. We propose such a procedure might be suitable for lip as well, and we suggest to systematically probe up to four orientations (0°, 45°, 90°, and 315°). For instance, this could be achieved by in a first step by determining the aMT for 45°, then for 0°, 90°, and 315°. As a second step, a series of MEPs could be collected at a supra-threshold intensity (e.g. ≥120% aMT) from each orientation to see which produces the largest responses that can be verified as MEPs based on the presence of a latency >7.5 ms and a silent period following the response.

In conclusion, our results for FDI and OO replicate and extend results of previous studies investigating optimal coil orientation in hand (Werhahn et al., 1994, Balslev et al., 2007). Our results indicate that coil orientations and associated induced current directions previously reported for muscles of the corticospinal tract, particularly FDI, were also appropriate for OO, a muscle innervated by the corticobulbar tract. However, the analysis pointed towards more variability in optimal orientation across participants with respect to the optimal coil orientation for eliciting the largest MEPs, so we recommend examining a range of coil orientations spaced between 315° and 90°Contralaterally.

